# Morning endurance training induces superior performance adaptations compared to afternoon training in mice

**DOI:** 10.1101/2023.09.18.557933

**Authors:** Stuart J. Hesketh, Collin M. Douglas, Xiping Zhang, Christopher A. Wolff, Casey L. Sexton, Elizabeth S. Nowicki, Karyn A. Esser

## Abstract

Endurance performance exhibits time-of-day variation in both humans and rodents, peaking in the late active-phase. However, whether the timing of endurance training influences performance adaptations remains unclear.

To investigate, female mice were trained 5-d/week for 6-weeks at either ZT13 or ZT22, using treadmill running at 70% of each animal’s maximal capacity. Endurance performance was assessed at baseline, week-3, and week-6. Secondary outcomes included blood glucose and lactate, cage activity, body composition, liver and skeletal muscle glycogen content, mitochondrial and contractile protein expression.

At baseline, late-active phase (ZT22)-tested mice exhibited significantly higher endurance capacity than early-active phase (ZT13)-tested mice (P<0.05). Following 6 weeks of training, ZT13-trained mice demonstrated a greater rate of improvement, with endurance increasing by 132% (P<0.05), compared to 45% in afternoon ZT22-trained mice. By week 6, performance improved but was similar between groups (P>0.05), despite lower absolute training volumes in the ZT13 group. Both training groups reduced fat-mass (ZT13: −31%,ZT22: −32%; P<0.05 vs. control), with no differences in lean mass, food intake or muscle and liver glycogen content (P>0.05). In skeletal muscle, ZT13-trained mice were associated with increased (P<0.05) COXIV protein expression, citrate synthase activity, and shifts in MyHC isoform expression, without changes (P>0.05) in mitochondrial content.

ZT13-training elicited superior performance adaptations despite lower absolute workloads, indicating enhanced training efficiency. These findings identify exercise timing as a biologically relevant factor influencing endurance adaptation and variability in exercise responses.

**NEW & NOTEWORTHY:** This study demonstrates that endurance training in the early active phase induces greater performance adaptations than late active phase training in mice, resulting in overcoming diurnal differences in exercise performance, despite lower absolute training volumes. These findings reveal exercise timing influences training efficiency, likely via circadian regulation of skeletal muscle metabolism. This work identifies time-of-day as a biologically relevant and underappreciated variable contributing to the heterogeneity of exercise responses, even in tightly controlled preclinical models.

## INTRODUCTION

Exercise is a remarkable stimulus that can profoundly affect whole body physiology and specifically alter skeletal muscle phenotype. Each coordinated exercise bout elicits an integrative systemic response to the subsequent increase in metabolic demands, driving chronic adaptations across multiple tissues (1, 2). Exercise capacity, therefore, relies on the orchestration of both physiological and metabolic responses among various tissues (1). Many of the metabolic health benefits associated with exercise center around skeletal muscle adaptations; for example, skeletal muscle is considered the largest metabolic tissue and a critical site for glucose disposal both at rest and during exercise (3, 4).

Endurance exercise performance varies by time-of-day in both humans and rodents, with better performance typically observed in the afternoon or later in the active period (5–9). In mice, these differences are dependent on a functional circadian clock. For example, genetic disruption of clock genes eliminates time-of-day differences in endurance capacity (7, 10). Previously, Adamovich et al., (7) have linked performance differences to liver glycogen availability, which is known to fluctuate with feeding and circadian rhythms. In this study, they implemented time-of-day–restricted exercise and found that the differences in performance were still evident after two weeks of training suggesting that timing may have less impact with adaptation. However, two-weeks is a very short duration for an exercise training study and longer-term time-of-day training studies, are lacking.

The circadian clock mechanism is an evolutionarily conserved transcription translational feedback mechanism that exists in virtually all cells, allowing organisms to align physiological functions with the 24-hour day. The core clock operates through a feedback loop of gene expression, where transcription factors BMAL1 and CLOCK regulate expression of genes *Period1/2* and *Cryptochrome1/2* which then feedback to inhibit their own expression and reset the cycle (11–15). While this molecular mechanism is conserved across tissues, the functional outputs of the clock are tissue-specific and critically shape physiology throughout the day. In skeletal muscle, for example, circadian rhythms influence maximum isometric strength and mitochondrial function, with peak performance and metabolic activity occurring at specific times of day (9, 16, 17). Disruption of the clock, either environmentally or genetically, impairs these time-of-day–dependent functions, as shown by reduced grip strength in mice lacking core clock components (18). Thus, understanding how the clock programs time-of-day variations in muscle function has important implications for optimizing health and performance.

Despite well-established time-of-day differences in acute exercise performance, relatively little is known about how the timing of exercise training influences long-term physiological adaptations and performance. Previous studies suggest that exercise can act as a non-photic time cue, capable of shifting the phase of peripheral clocks, particularly in skeletal muscle (19–22). However, it remains uncertain whether training at specific times of day enhances or impairs adaptation or performance. Understanding whether consistent training during the early versus late active period differentially impacts endurance capacity, metabolic outcomes, or body composition, and phenotype is not only critical for understanding the influence of exercise-timing has on the heterogeneity of the exercise response, but for optimizing exercise prescriptions in both health and disease contexts.

Herein, we investigated how the timing of endurance training influences performance and muscle tissue adaptations in mice. Using a 6-week treadmill training protocol, we compared mice trained at the beginning (ZT13) or end (ZT22) of their active phase at workloads matched for relative exercise intensity. We assessed maximal endurance performance before training, mid-training (3wk) and post-training (6wk) and evaluated secondary outcomes, including blood glucose and lactate responses, voluntary cage activity, body composition, and muscle biochemistry. Our primary objective was to determine whether consistent training in the early active phase versus the late active phase confers differential performance adaptations with selected targeted physiological and metabolic outcomes. Our results demonstrate that mice trained in the early active phase exhibited greater performance adaptations compared to those trained in the late active phase. These differences in training efficiency resulted in the mice trained in the early active phase achieving the same maximum endurance performance as those in the late active phase by 6 weeks of training. It was interesting to note that this was not evident until after 3 weeks of training, suggesting that the tissue and systemic adaptations to support improved performance required a significant duration of training to be realized. We did not detect differences between exercise groups in resting levels of muscle or liver glycogen after training; however, markers of skeletal muscle oxidative metabolism and modest shifts in contractile protein expression were observed in ZT13 compared with ZT22 runners. We posit that time-of-day-dependent adaptations to endurance run training may act directly through the molecular clock mechanism in muscle and potentially other organ systems to support differential efficiency of performance adaptations.

## MATERIALS AND METHODS

### Ethical approval

All animal procedures in this study were conducted in accordance with the guidelines of the University of Florida for the care and use of laboratory animals (IACUC #201809136). The use of animals for exercise protocols was in accordance with guidelines established by the US Public Health Service Policy on Humane Care and Use of Laboratory Animals.

### Animals

Eighteen female PERIOD2::LUCIFERASE (PER2::LUC) mice on a C57/Bl6J background (23) aged 5 months (22 ± 1 g body weight) were bred in-house from mice originally received as a gift from Dr. Joseph Takahashi. Previous data from our laboratory reported no sex-specific effects of exercise training on muscle circadian PER2::LUC phase (19), so female mice were selected, as they are known to run more than their male counterparts (24). Mice were initially housed (12hr light:12hr darkness; ZT0 = time of lights on/ rest phase, ZT12 = time of lights off/ active phase) in a controlled climate (23 ± 1.5 °C, 58 ± 5.8 % relative humidity) and had *ad libitum* access to water and standard rodent chow (Envigo Teklad 2918, Indianapolis, IN, USA). Mice were then moved to single housing with the same light, climate, and nutritional conditions prior to starting the experimental protocol. All experiments took place during the dark/ active phase, and all treadmill testing and training took place in the dark, under red light, on a Panlab treadmill (Harvard Apparatus, Holliston, MA). All mice were anaesthetized using isoflurane and euthanized by cervical dislocation under red light, ∼3 days after the last bout of exercise training at ZT17. All tissues analysed were collected at the same time-of-day and included the gastrocnemius and liver which were isolated and cleaned of fat and connective tissue then frozen in liquid nitrogen and stored at −80°C pending further analysis. We collected tissues for real-time PER:LUC bioluminescence data capture but unforeseen circumstances resulted in these data being lost prior to submission.

### Maximal Endurance Capacity Testing

Each maximal endurance capacity testing session was performed, similar to that described in Maier et al., (5). Briefly, animals began at a speed of 10 cm/s at 10° incline for a 5 min warm-up period. The incline was increased to 15° and the treadmill speed was increased by 3 cm/s every 2 min until exhaustion. The treadmill was operated with the electrical shock grid turned off and sponges were placed at the back of the treadmill to reduce the risk of injury. Mice were deemed exhausted when they remained in contact with the sponge >10 s and could not be encouraged to continue by several air puffs from a compressed air container. All mice were tested at the time which corresponded to their group (i.e., either ZT13 or ZT22). The sedentary control group (n = 6) were split so that half (n = 3) were handled at ZT13 in the same manner as the early active phase training group, and the other half were handled at ZT22 identically to the late active phase training group. This involved being moved from the housing suite into the treadmill room for the duration of each exercise session, here they maintained a sedentary state in their cages and were positioned next to the treadmill.

### Maximal Endurance Testing Schedule

This study is focused on time of training and the training times used were a) the early active phase which is defined by occurring at 1hr after lights off in the animal facility (ZT13) and b) the late active phase which is defined by occurring at 10hrs after lights off (ZT22). Initially, mice were randomized into two groups for pilot maximal endurance capacity testing, an early active phase group (ZT13, n = 9) and a late active phase group (ZT22, n = 9). This was to confirm that there were measurable time-of-day differences in exercise capacity which were consistent with prior reports (6, 25). Immediately prior to the pre-training trial, mice were subjected to three days of treadmill familiarization as in previously described methods (25). Briefly, during the first day of familiarization the speed of the treadmill was set to 10 cm/s for 5 min at 0° incline. The incline was then adjusted to 5° and speed was ramped up by 2 cm/s every 2 min up to 20 cm/s. The second day consisted of an increase in speed every 3 min by 3 cm/s from 10 cm/s to 24 cm/s at an incline of 10°. The final familiarization session was the same as the second day but performed on a 15° incline. On the day after the third familiarization session, maximal endurance capacity testing was performed. This was followed by a 10 day washout period where mice were acclimated to new housing. Mice were continuously monitored for daily cage activity using wireless, infrared activity monitoring (Actimetrics, Wilmette, IL, USA; analyzed using ClockLab software). Mice were then re-randomized into three experimental groups where they remained for the 6-week time-of-day training: i) training at ZT13 (n = 6), ii) training at ZT22 (n = 6), or iii) sedentary control (CON, n = 6). These groupings denoted the time-of-day in which testing and training occurred. Similar to the pilot testing the CON group was split so that half (n = 3) were handled at the same time as the early active phase group, and the other half were handled with the late active phase group. This involved being moved from the housing suite into the treadmill room for the duration of each exercise session, where they were maintained sedentary in their cages and positioned next to the treadmill. Mice had their food removed 1 hour prior to exercise testing and were assessed for their maximal endurance capacity before, after 3 weeks, and after 6 weeks of time-of-day training. Endurance capacity testing conducted after 3 weeks was used to scale exercise intensity to account for training adaptations and assess a time course for improvements. A schematic of the study design is shown in Figure 1.

**Figure 1.**
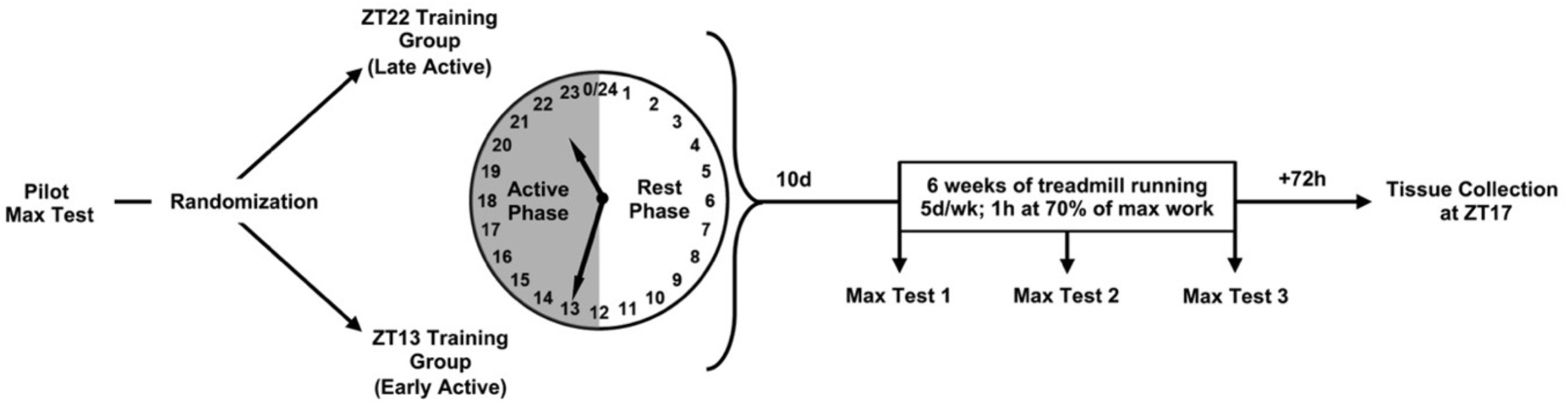
Study Design. A schematic of the experimental design is presented to show the time-line of testing, grouping, and training. Mice were randomized for pilot testing, re-randomized into their experimental training groups ZT13 (early active phase runners) or ZT22 (late active runners), then 10 days later mice were re-tested (Max Test 1), and the 6 weeks of training began. d = days, h = hours, wk = week, ZT = zeitgeber time.

### Endurance Training Program

The training program consisted of 5 exercise bouts per week, over the 6 weeks of training (30 individual exercise bouts), all bouts were consistently performed at the assigned time-of-day training group times (i.e., ZT13 or ZT22). Each training bout consisted of 1 hour of treadmill running at a consistent slope of 15° and a speed corresponding to 70 % of work done during maximal endurance testing, calculated similar to equations from Avila et al., (26) and shown in equation 1 below.

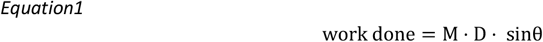

Were M is the mass (g) of the animal, D is the distance (m) ran in the maximal capacity test and θ is the slope (°) of the treadmill. Work done is measured in arbitrary units (AU).

### Blood collection and body composition

Tail blood glucose (Accu-Chek, Roche) and blood lactate (Lactate Plus meter, Nova Biomedical) values were determined immediately prior to the maximal exercise capacity test and immediately post, within 1 min after physical exhaustion. Blood glucose and lactate was also taken on the third session of each week both pre and post. Body composition was assessed by nuclear magnetic resonance (Echo MRI-100 Body Composition Analyzer; Echo Medical Systems, Houston, TX) and was conducted in the middle of the active phase at ZT17, on the days immediately prior to maximal testing.

### Muscle and liver glycogen

Small (∼10 mg) frozen pieces from the gastrocnemius muscle and liver were used to detect tissue glycogen content. Care was taken to ensure the same location of each tissue was sampled across replicates. Tissue samples were weighed and homogenized in 100 μl of glycogen hydrolysis buffer in a bullet blender (BBY24M, Next Advance, NY, USA). Homogenates were centrifuged at 12000 rpm for 5 mins at 4 °C. Muscle glycogen concentrations were then quantified from the supernatant using a commercially available kit (K2144, ApexBio, Houston TX, USA) according to manufacturer’s instructions. Glycogen concentrations were plotted against a standard curve and background glucose was subtracted to calculate glycogen content which was expressed normalized to tissue weight.

### Western blot

Muscle lysates were prepared using cryo-pulverized samples from the quadriceps muscle group. Cryo-pulverization was used instead of selected tissue sampling to minimize potential for regional variation from this large muscle group. Samples were prepared for western blots and protein loading ranged from 4 ug/sample for MyHC gels to 20 ug protein/sample for mitochondrial proteins. Our SDS-PAGE gels for protein separation were either 7 % or myosin heavy chain and citrate synthase proteins or 10 % gels for the smaller proteins. Proteins were transferred to PVDF membranes following standard procedures in the lab (27) and following transfer, membranes were blocked with blocking buffer (5 % non-fat dry milk in 0.1 % TBS-T; Blotto, Santa Cruz Biotechnology sc-2324) for 1 hour at room temperature. Following blocking, membranes were incubated overnight at 4°C with primary antibodies specific to Type I, IIa, IIb, and IIx MyHC, citrate synthase, VDAC, COX IV, and gamma-tubulin (Developmental Studies Hybridoma Bank antibodies BA-D5 Type I MyHC 1:500, SC-71 Type IIa MyHC, 1:500 BF-F3 Type IIb, MyHC 1:1000, Invitrogen PA5-31466 1:1000; Invitrogen PA5-22126 1:2000; Cell Signaling 4661 1:1000; Cell Signaling 4850 1:1000; Millipore Sigma Anti-gamma-tubulin antibody T6557) in 1:1 diluted blocking buffer with 0.1 % TBS-T before incubation with secondary HRP conjugated secondaries for 1 hour at room temperature in diluted blocking buffer. Following secondary antibody incubation, membranes were washed 3×5 minutes with 0.1 % TBS-T and 1×5 minutes with ultrapure water, prior to incubation with ECL reagent followed by imaging.

### Citrate Synthase Activity

Entire quadriceps muscle was cryo-pulverized and mixed over liquid nitrogen to obtain a homogenous sample, and a ∼20 mg aliquot was taken for analysis of citrate synthase activity. Cryo-pulverized tissue was homogenized in 20 volumes (weight/volume) of buffer consisting of 20mM Tris-HCl, 137mM NaCl, 1% Triton X-100, 2mM EDTA, and protease inhibitor cocktail (mini-cOmplete EDTA-free, Roche). Protein concentrations of homogenized samples were obtained using the RCDC Protein Assay (Bio-Rad), performed to the manufacturer’s specifications. Citrate synthase activity was then obtained using the MitoCheck^®^ Citrate Synthase Activity Assay Kit (Cayman Chemical), performed to the manufacturer’s specifications. In brief, 30 uL of homogenate was used as input, and absorbance was recorded per manufacturer’s specifications via plate reader set at 412 nm and 25°C for 10 min. Citrate synthase activity was then calculated per provided formula using protein concentrations of skeletal muscle homogenates, and plotted as nmol/min/mg of skeletal muscle tissue.

### Statistical analysis

Unless stated otherwise, data are presented as mean ± standard deviation and all statistical analyses were conducted in GraphPad Prism 9.1.2 (GraphPad Prism, RRID:SCR_002798). For multiple comparisons, data were analyzed using one or two factor analysis of variance (ANOVA) followed by Tukey’s multiple comparisons test. To assess differences between exercise training groups only, two-tailed independent student t-tests were performed to evaluate statistical differences. All statistically significant thresholds were considered at the level of P<0.05. All raw data that support the findings of this study are available from the corresponding author upon reasonable request.

## RESULTS

### Early active phase runners exhibited enhanced adaptations compared to late active phase runners

To evaluate the effects of time-of-day treadmill training on maximal endurance performance, we first performed a maximal endurance capacity test prior to training at two distinct times during the active period. In accordance with previous work in humans (28) and mice (6). We established that mice tested in the late active period: ZT22 exhibited significantly greater treadmill endurance capacities than those tested in the early active period: ZT13 (Figure 2A). To allow for comparisons among the mice, we calculated treadmill work done which considers the treadmill speed, incline, duration, and individual mouse bodyweight. We determined that the late active phase runners completed 85% more treadmill work than the early active phase runners (2552 ± 260 arbitrary units (AU) vs. 1381 ± 354 AU) (P<0.001; Figure 2A). Additionally, mice tested in the ZT22 achieved an 83% further distance (P<0.001; Figure S1A) and spent 57% longer duration on the treadmill (P< 0.001; Figure S1B) than their ZT13 testing counterparts.

**Figure 2.**
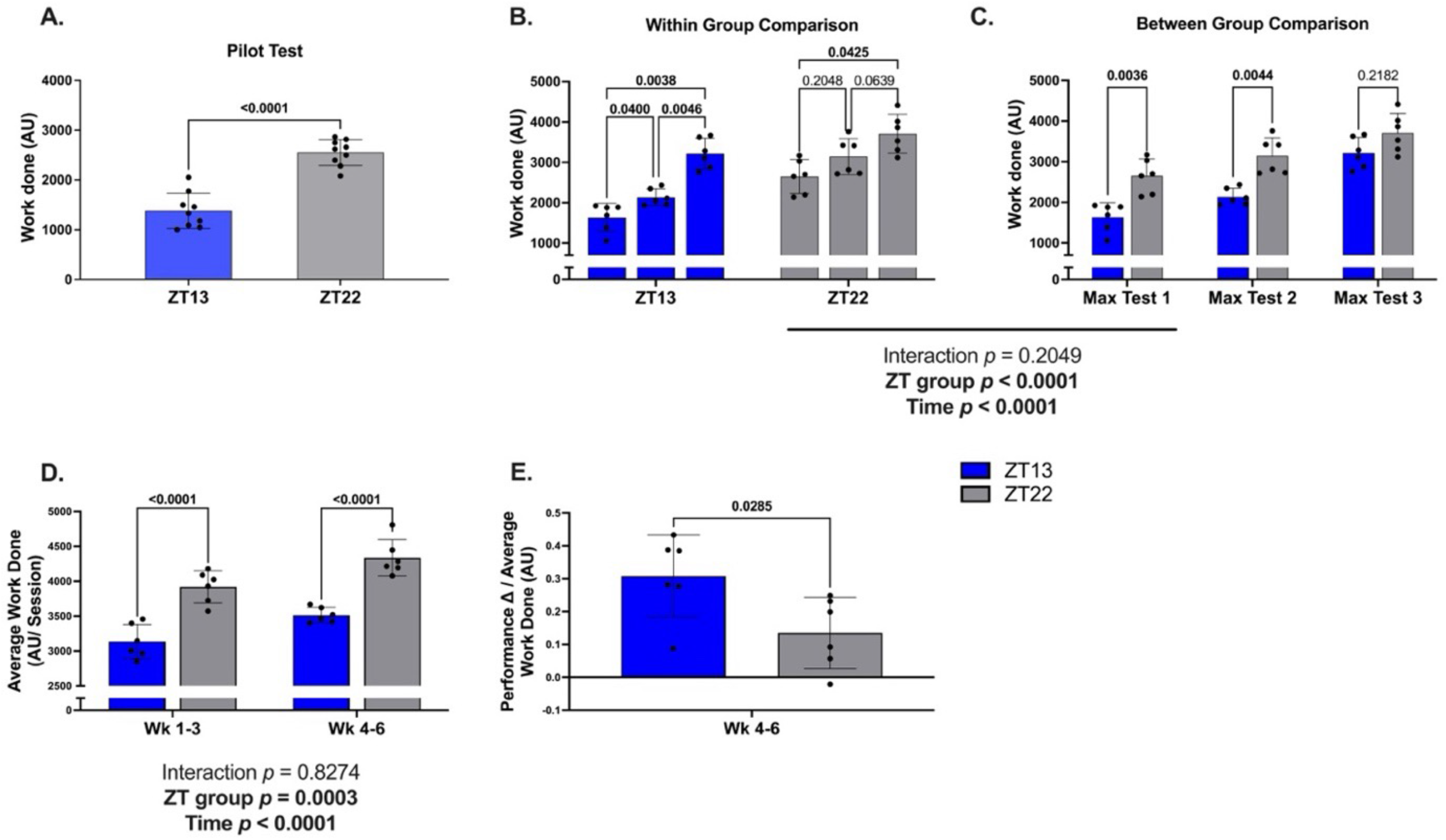
Time of exercise training reduces diurnal differences in exercise performance. All data are presented as MEAN ± SD with individual values plotted, and significant P-values are bolded in the figure for identification. Each maximal test was conducted at ZT13 (blue bars) for the early active phase runners and at ZT22 (gray bars) for the late active phase runners. **A**) Ten days prior to the onset of training, a pilot test was conducted to confirm if diurnal differences in maximal run performance existed for early active phase runners (n = 9) and late active phase runners (n = 9). **B, C)** Maximal endurance capacity testing was conducted prior to (Max test 1), 3 weeks after (Max test 2), and 6 weeks after (Max test 3) the onset of training (ZT13 runners: n = 6; ZT22 runners: n = 6). **D)** The average work done per training session for ZT13 and ZT22 runners was compared across weeks 1-3 and weeks 4-6. **E)** Change in run performance between Max Test 2 and Max Test 3, divided by the average work done per session from week 4 to 6 was compared between ZT13 and ZT22 runners. One and two factor analysis of variance were used to assess statistical significance, apart from panels A and E were two-tailed independent T-tests were performed. Post-hoc comparisons of ZT13 and ZT22 runners are also shown through within-group and between-group comparisons. All statistically significant thresholds were considered at the level of P<0.05.

Ten days following pre-testing, mice were randomly assigned to complete 6 weeks of scheduled run training during the early active phase or late active phase. Maximum endurance capacity was tested at 3 timepoints; i) prior to the onset of training, ii) after 3 weeks of training, and iii) after 6 weeks of training. Within each time-of-training group, we found that the ZT13 mice exhibited significant increases in endurance capacity from onset to 3 weeks (P= 0.040) and 3 weeks to 6 weeks of training (P= 0.005). In contrast, the ZT22 training group demonstrated a significant change in endurance performance when comparing between the pre-test to 6 weeks (P= 0.043), but endurance capacity at 3 weeks was not statistically different from either onset (P= 0.205) or week 6 values (P= 0.064) (Figure 2B). The plotted trajectory of individual mouse endurance time is provided in supplemental data (Figure S2) which suggests that rate of change in endurance capacity is greater in the early active phase trained compared to the late active phase trained mice.

In Figure 2C we compared maximum endurance capacity (work done) between each time-of-training group for each maximal test. After 3 weeks of training, the ZT22 mice continued to perform significantly better, completing more treadmill work (47%; P= 0.004) compared to those tested at ZT13. However, following 6 weeks of run training, the endurance performance of mice that trained during the early active phase increased and was not different from those that trained during the late active phase (P= 0.218) (ZT13 runners: 3214 ± 384 AU vs ZT22 runners: 3708 ± 481 AU; Figure 2C). Together, these endurance performance results indicate that mice in the early active phase training group exhibited a greater rate of adaptation in endurance performance compared to the mice trained in the late active phase. By week 6, ZT13 runners exhibited no difference in treadmill performance compared to the ZT22 runners.

Lastly, since both groups of mice were trained at the same relative training intensity, we wanted to examine the differences in absolute work done during training between early active phase runners and late active phase runners to understand how differences in the average amount of total work done may have affected maximal performance adaptations (Figure 2D). We show work done on average across weeks 1 to 3 (P< 0.001) and weeks 4 to 6 (P< 0.001) of training was significantly greater for ZT22 runners compared to ZT13 runners. Given that ZT13 and ZT22 runners showed no difference in maximal endurance capacity at max test 3, but ZT13 runners did less work on average across weeks 4-6, we asked if the ZT13 runners experienced significantly outsized adaptations to training (Figure 2E). We calculated the change in performance between week 3 (Max test 2) and week 6 (Max test 3) to assess adaptation during the second half of the training program, and discovered that the change in performance per average work done across weeks 4 to 6 was significantly greater (P= 0.029) for ZT13 runners compared to ZT22 runners. Therefore, ZT13 runners were subjected to less absolute work, while training at equal relative workloads. This means that runners in the early active phase demonstrated greater adaptation per the amount of work completed during training, i.e. enhanced efficiency of performance adaptations.

### Blood glucose, blood lactate response to exercise and tissue glycogen content with training show no time-of-day differences

To begin to try and identify potential markers of time-of-day exercise capacity outcomes, we tested well-established markers of acute exercise and training responses. To interrogate the acute response during maximal endurance exercise we measured blood glucose and blood lactate immediately before and after each maximum endurance test. We observed significantly higher measures of blood glucose (P< 0.001) and blood lactate (P< 0.001) immediately after each maximum capacity test (Figure 3A & 3B). However, there was no difference (P>0.05) between ZT13 and ZT22 runners for either measure, and there was no change (P>0.05) with training. Thus, the acute glucose and/or lactate response to a maximal graded exercise test does not differ between time-of-training groups and does not show any pattern that associates with differences during any endurance capacity tests.

**Figure 3.**
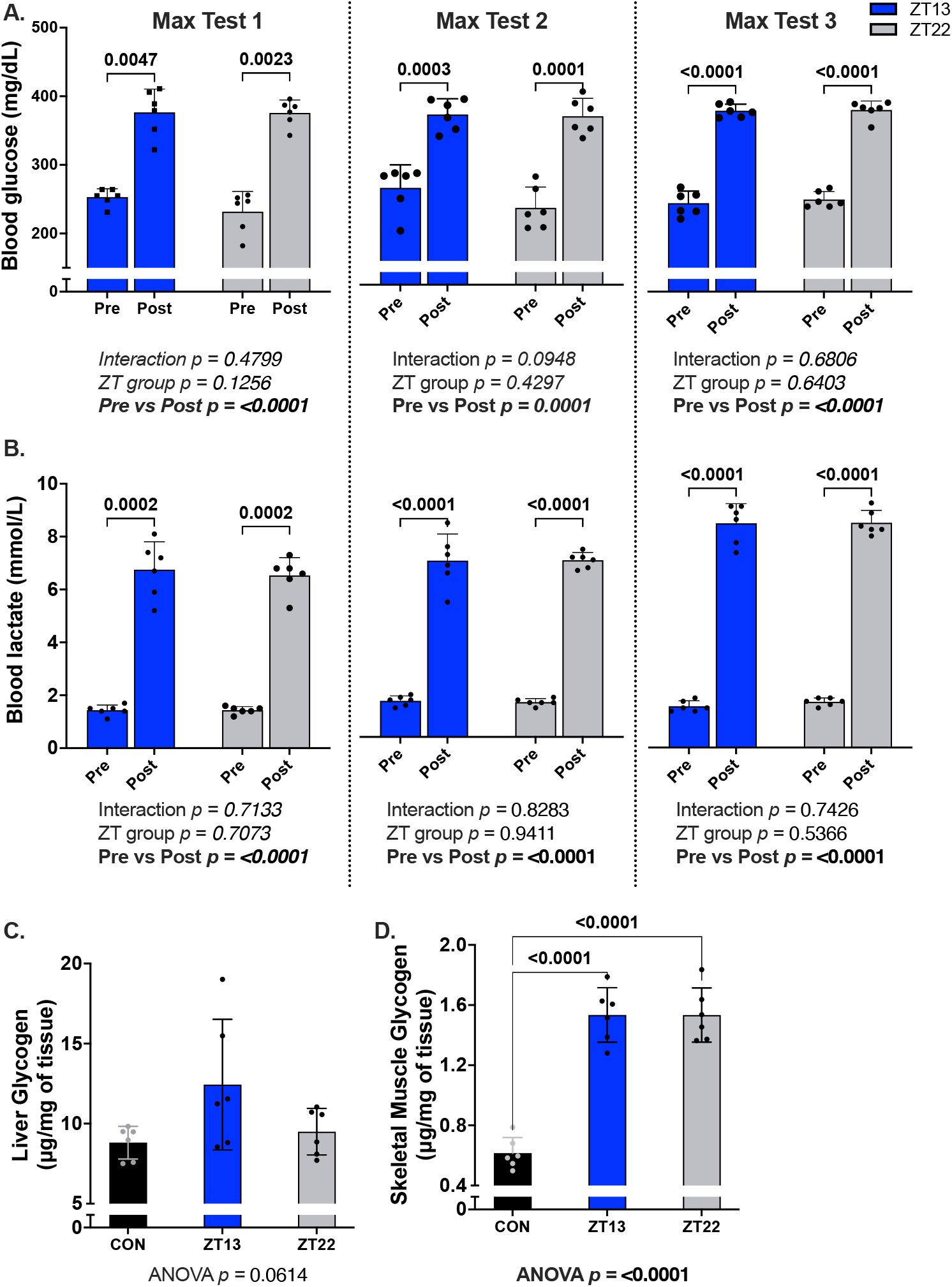
Maximal capacity test blood markers and basal tissue glycogen content. All data are presented as MEAN ± SD with individual values plotted, and significant P-values are bolded for identification. Each maximal test in A and B was conducted at ZT13 for the early active phase runners (blue bars), (n =6), and at ZT22 for the late active phase runners (gray bars), (n = 6). **A, B)** Acute blood markers of glucose and lactate were collected both pre and post each maximal testing session. **C, D)** Show the tissues used for glycogen content analysis, liver (µg/mg of tissue) and skeletal muscle (µg/mg of tissue), samples collected 72h after the last exercise bout. Two-tailed independent T-tests were performed to assess statistical significance between pre and post measures in panel A and B, and two-way analysis of variance were used to assess interaction. One way analysis of variance and Tukey’s multiple comparisons test were used to assess between group differences in panel C-D. Statistically significant thresholds were considered at the level of P<0.05 for all.

Muscle and liver glycogen levels are also well associated with exercise training and performance (7, 29, 30). Consistent with sampling strategies performed in the MoTrPAC inititive (31), we performed assays for glycogen on muscle and liver tissues collected 72 hours after the last bout of exercise at a common timepoint midway between the time of early active phase and late active phase training, ZT17. It is well known that endurance training results in increased glycogen storage in both muscle and liver (32, 33). After 6 weeks of training, we determined that skeletal muscle increased glycogen content in both ZT13 and ZT22 training groups but that there was no difference (P> 0.05) between the groups. We also determined that liver glycogen levels at ZT17 were not different with exercise training between groups and these levels were not different compared to the sedentary controls. However, we did note, that there is a trend for a difference in liver glycogen across the groups (P= 0.061) but with the high variability in the liver glycogen levels for the early active phase exercise group we were underpowered to have confidence in the outcomes. Thus, we did not detect any time-of-day training effects on the magnitude of glycogen storage in muscle or liver.(34)

### Increased Daily Cage Activity and Body Composition during Exercise Training

We next asked if there were differences in the daily cage activity in the mice in the ZT13 or ZT22 training groups. For these experiments we used infrared motion sensors in the home cage to track 24 h movement around the cage. We found that mice from both training groups exhibited increased movement in their home cages compared to the control mice throughout the 6 weeks of training (Figure 4A). We noted that the activity of the mice was largely limited to the normal active period (dark phase) with no indication of altered activity in the light or rest phase of the day (Figure S4). We did detect cage activity differences between the early active phase and late active phase training groups with the late active phase runners showing more daily cage activity in the first week (P= 0.002), while the early active phase runners had more cage activity in the last week of training (P< 0.001). It is important to note that these measures are reflecting behavioral changes based on movement in the cage, but these measures do not provide distance travelled.

**Figure 4.**
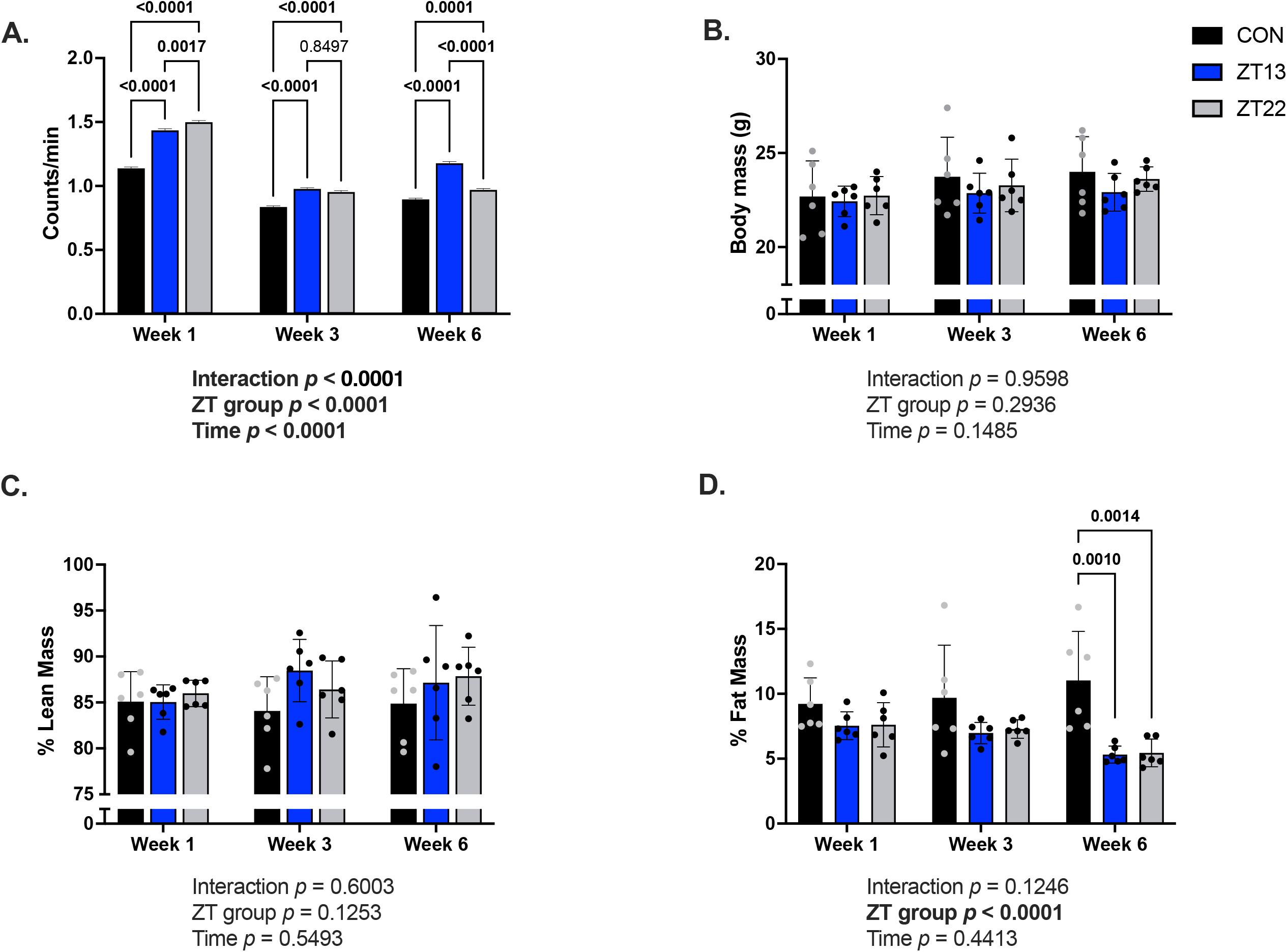
Mouse cage activity and body composition profiles. Unless stated otherwise, data are displayed as MEAN ± SD with individual values plotted, and significant P-values bolded for identification. Mice were individually housed for the duration of the time-of-day training program, and all animals were evaluated at weeks 1, 3, and 6 for cage activity and body composition. The sedentary control group is shown in black (n = 6), the ZT13 early active phase runners in blue bars (n =6), and the ZT22 late active phase runners are shown in grey bars (n = 6). **A)** Shows average data for each group for daily cage activity using wireless, infrared monitors in counts per minute. **B)** Displays total body mass over time. **C, D)** Show percent lean mass, and percent fat mass, respectively. Measured at ZT17 each week immediately prior to maximal testing using Echo MRI. One way analysis of variance and Tukey’s multiple comparisons test were used to assess between group differences and interaction. Statistically significant thresholds were considered at the level of P<0.05.

We also sought to investigate if time-of-training elicited differential changes in body mass and/or body composition and if there were any alterations that associated with the differences in work done during treadmill running. We measured weekly food consumption in all mice and found there were no significant differences in the amount of food consumed across all 3 groups (Figure S3). We tracked body weight and body composition at weeks 1, 3, and 6 following training in all groups. There were no significant differences between sedentary control, early active phase runners, or late active phase runners in body mass (Figure 4B) or percent lean mass (Figure 4C). However, percent fat mass was significantly reduced at 6 weeks in both early active phase (P= 0.001) and late active phase runners (P= 0.001) compared to sedentary controls (Figure 4D). These results indicate that time-of-day training over 6-weeks did not significantly alter body mass or lean mass but did reduce fat mass in both training groups. Because there are no differences between time-of-day training groups, we suggest changes in fat mass do not correlate with the time of training performance differences. However, it is interesting to note that the ZT13 runners did less absolute physical work than the ZT22 runners yet the fat mass reduction with training was not different.

### Muscle mitchondrial enzyme activity and protein expression and myosin heavy chain isoform expression following time-of-day training

Increased citrate synthase activity has been well studied as an adaptation to endurance training (34), often being utilized as a marker for aerobic capacity (35) and mitochondrial volume density (36). Therefore, we measured citrate synthase protein content and enzyme activity in quadriceps muscle homogenates to ask if there are differences in the muscle of early active phase or late active phase runners (Figure 5H-I). Citrate synthase protein levels showed no difference (p > 0.05) between any of the groups (Figure 5H). However, consistent with the literature for citrate synthase activity (Figure 5I) analysis identified a significant difference (P= 0.001) between the control group (8.159 ± 0.693 nmol/min/mg), and both trained groups; ZT13 runners (10.810 ± 0.439 nmol/min/mg), and ZT22 runners (6.339 ± 1.729 nmol/min/mg). However, there were no differences in citrate synthase activity between the trained groups.

**Figure 5.**
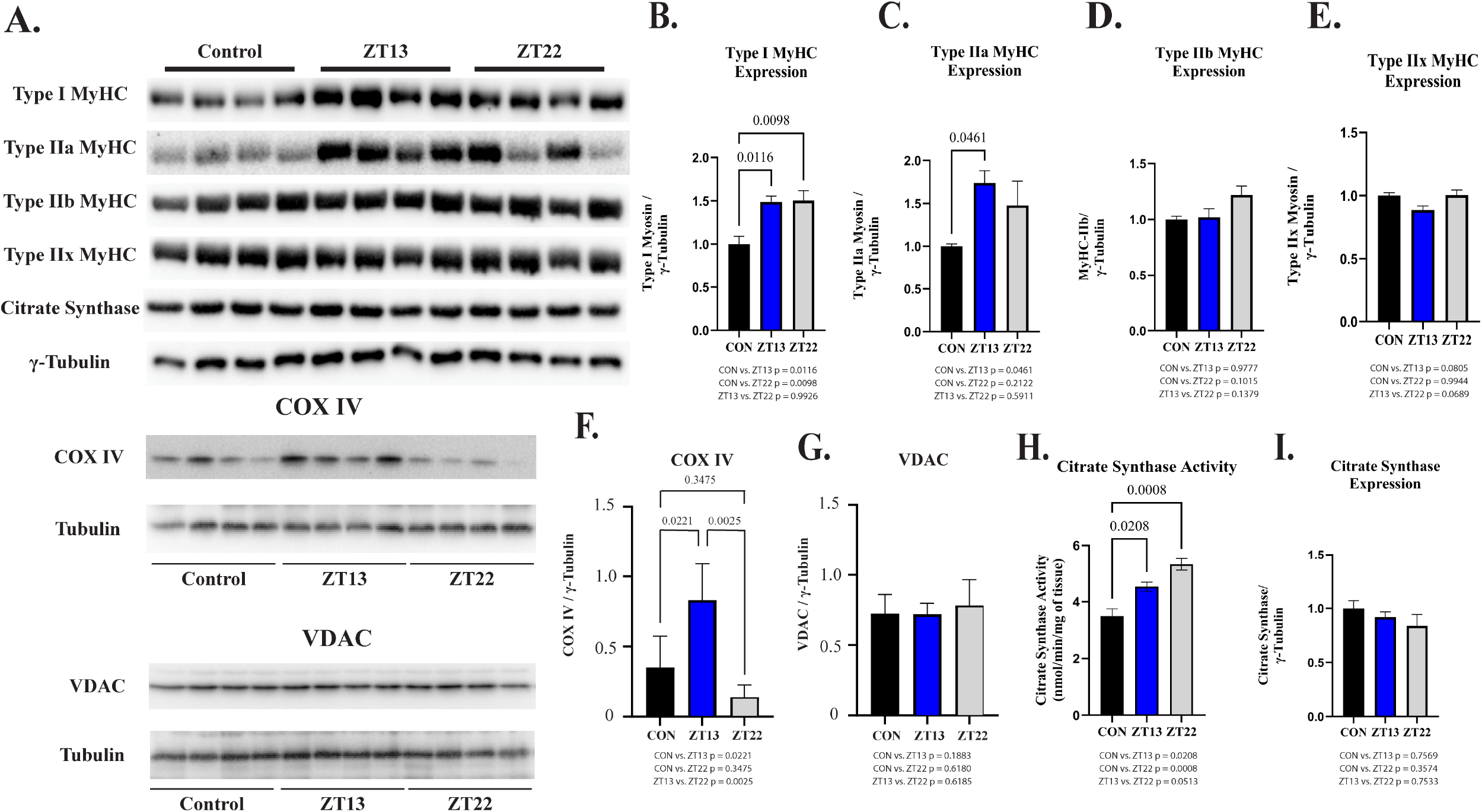
Time-of-day specific effects of endurance training on mitochondrial and myofibrillar protein expression in quadriceps muscle. Data are presented as mean ± SEM. Control (CON, n = 4), early active phase trained (ZT13, n = 4), and late active phase trained (ZT22, n = 4) groups are shown. Effects of early and late active phase endurance training on mitochondrial and myofibrillar protein expression in quadriceps muscle. Quadriceps muscles were collected 72 h after the final training session, cryopulverized, and analyzed by immunoblotting. **A)** Representative immunoblots for myosin heavy chain (MyHC) isoforms (Type I, Type IIa, Type IIb, and total MyHC), citrate synthase, COX IV, and VDAC, with γ-tubulin used as a loading control. Data plotted is relative to control average. **B–E)** Quantification of Type I, Type IIa, Type IIb, and total MyHC protein abundance, normalized to γ-tubulin and expressed relative to non-exercised control animals. Data plotted is relative to control average. **F)** COX IV protein abundance normalized to γ-tubulin. **G)** VDAC protein abundance normalized to γ-tubulin. **H)** Quantification of citrate synthase activity. **I)** Quantificataion of citrate synthase protein normalized to γ-tubulin. Data plotted is relative to control average. Statistical comparisons were performed using one-way ANOVA with Tukey’s post hoc test. *P* < 0.05, P < 0.01; absent comparisons were not statistically significant.

To further explore potential mechanisms underlying the superior performance adaptations in the early active phase group, we examined mitochondrial and myosin heavy chain isoform protein expression in quadriceps muscle homogenates. Cytochrome c Oxidase Subunit IV (COX IV) protein abundance, a key component of complex IV (Figure 5F) was significantly higher in early active phase trained mice compared with controls (p < 0.05) and was not different (p > 0.05) in the late active phase group, whereas Voltage-Dependent Anion Channel (VDAC) content (Figure 5G) which is a protein in the mitochondria outer membrane did not differ between any of the groups. Analysis of myosin heavy chain isoforms found that between exercise groups there were no significant differences observed in Type I, IIa, IIb or IIX myosin heavy chain (MyHC) isoform expression (Figure 5B-E). However, greater (p > 0.05) levels of Type I and IIa MyHC were detected in ZT13/early runners, but not in ZT22/late runners Type I MyHC was the only elevated isoform when compared to non-exercised control animals. Together, these findings suggest that large-scale changes in content of mitochondria or composition of myosin heavy chain isoforms are unlikely to fully explain the observed performance differences. Rather, the COXIV data are consistent with changes in key components of metabolic capacity that may preferentially occur with early active phase training.

## DISCUSSION

Our study provides novel insight into the effects of exercise timing on long-term performance and physiological adaptations to endurance training in mice. We have demonstrated that while mice initially exhibited superior endurance exercise performance in the late active phase, consistent early active phase training over six weeks resulted in greater improvements in endurance capacity compared to late active phase training (Figure 2E). Remarkably, these adaptations occurred despite both groups training at the same relative intensity, so a greater magnitude of performance was attained by the ZT13 group despite performing less absolute treadmill work. This provides evidence suggesting that time of exercise is a modifiable aspect of a training program focused on enhanced endurance performance outcomes. Of note, we also found that the ZT13 runners lost similar fat mass to ZT22 runners with no change in lean mass (Figure 4D) even though the absolute workload was less, suggesting differential reliance on exercise substrates linked to time of exercise. Lastly, we identified the mitochondrial marker, COX IV protein (Figure 5F), as a potential factor linking the early active phase training to the enhanced performance efficiency.

Cytochrome c oxidase subunit IV (COX IV) was selected for analysis as a functionally relevant and dynamically regulated component of mitochondrial complex IV. COX IV is a nuclear-encoded subunit that modulates electron flux through the catalytic core of complex IV and plays a critical role in matching mitochondira respiration to cellular energy demand via ATP- and ADP-dependent regulation. Importantly, prior studies have demonstrated time-of-day variation in maximal oxygen consumption and mitochondrial oxidative capacity in both humans (9) and rodents (37), implicating circadian regulation of electron transport chain function. Moreover, circadian oscillations in mitochondrial protein abundance, including components of oxidative phosphorylation complexes, have been reported in mouse liver (38), consistent with clock-controlled regulation at the protein level rather than through large changes in mitochondrial content. COX IV is also characterized by a relatively short protein half-life, making it particularly amenable to circadian modulation. In support of this concept, rhythmic variation in complex IV activity has been demonstrated in Drosophila (39), further suggesting that complex IV represents a node through which circadian timing can influence mitochondrial efficiency.

In the present study, COX IV protein abundance was selectively increased in early active phase (ZT13) trained mice despite no detactable differences in VDAC abundance (Figure 5), indicating that early active phase training may promote qualitative, enzyme-level enhancements in mitochondrial oxidative capacity rather than increased mitochondrial content. Notably, COX IV was assessed 72 hours after the final training session indicating that the increase in COX IV reflected a steady state change with early active phase training. However, our analysis was only done at one time of day, ZT17 and we cannot determine if COX IV protein levels are changing over time of day. If protein levels do change over time of day, the difference between the early vs. late active phase runners may reflect changes in circadian clock setting leading to a different temporal pattern of peak COX VI protein. Future studies incorporating longitudinal sampling across training and across time of day will be necessary to establish the temporal relationship between mitocondrial enzyme regulation and performance adaptation.

Western blot analyses of quadriceps muscle did not reveal statistically significant differences in overall myosin heavy chain (MyHC) isoform composition between early and late active phase training groups (Figure 5). However, differences were evident when each group was compared with non-exercised controls. Specifically, Type I MyHC protein abundance was increased in both early and late active phase trained mice, whereas a modest increase in Type IIa MyHC was observed only in the early active phase group. Interpretation of these findings requires consideration of the quadriceps muscle group, which in mice is predominantly composed of fast, glycolytic fibres (Type IIB and IIX), with more oxidative fibres localized to specific regions such as the rectus femoris and vastus intermedius. Thus, small changes in oxidative MyHC isoforms detected in whole-muscle homogenates likely reflect subtle, regionally constrained adaptations rather than large-scale fibre-type remodeling. These observations are consistent with prior work (40, 41) demonstrating that endurance training can induce gradual and localized increases in oxidative MyHC expression without necessitating overt fibre-type transitions, particularly when assessed at the whole-muscle level. Collectively, these data suggest that early active phase training may favor modest shifts toward a more oxidative contractile phenotype that complement mitochondrial adaptations, potentially contributing to enhanced training efficiency without major structural remodeling.

(38)Endurance exercise capacity in mice has been shown by our lab and others to be lower during the hours of the early active phase compared to the late active phase hours (5–7, 25). In this study, we used genetically similar mice and carefully controlled for age, sex and, relative exercise training intensity to better isolate the effect of training time on adaptation. Despite the initial performance disadvantage, ZT13-trained mice, exhibited a steeper trajectory of performance improvement compared to ZT22-trained mice (Figure 2E), ultimately achieving equivalent performances by week six (Figure 2C). This greater adaptation per unit of work means that the early active phase runners exhibited enhanced efficiency with training to yield improved maximal performance outcomes. The observation of differential training efficiency has not previously been documented. We suggest that our use of tight controls over many parameters of this study were necessary to demonstrate the time-of-day training differences in mice.

Notably, these performance improvements in the ZT13-trained mice were not detectable after 3 weeks of training, indicating that overcoming time-of-day performance differences requires an extended training duration. Our results align with prior observations from Adamovich et al., (7), who reported persistent diurnal performance differences after 2 weeks of time-of-day specific run training in mice. In our study, only after 3 weeks, did the early active phase group close the diurnal performance gap. Our observations are further consistent with findings from Souissi and colleagues who demonstrated 6 weeks of resistance training in humans was sufficient to overcome time-of-day maximal strength differences, with effects persisting at 2 weeks post-training (42). Together, these data suggest that while early active phase training may be less optimal for acute performance, it can elicit robust long-term adaptations at a lower absolute workload, provided the training is sustained more than 3 weeks.

Our results raise an important question about the mechanism(s) that underly the enhanced endurance performance outcomes as well as the enhanced fat loss per amount of treadmill work found in the early active phase training mice. We hypothesize that changes in the circadian clock mechanisms in skeletal muscle contribute to enhanced metabolic shifts that support the increased efficiency of performance adaptation in the early active period runners. The evidence for this model starts with studies that link time of exercise, the muscle clock and mitochondrial function. Specifically, Wolff & Esser, (19) demonstrated that time of exercise training was sufficient to shift the steady state phase of skeletal muscle clocks ∼4hrs after 6 weeks of training in the light/inactive phase. Beyond keeping time, the circadian clock regulates a time-of-day pattern of gene expression, and several groups have demonstrated that mitochondrial structure and function vary over time-of-day with higher function in the late active phase (8, 9, 43–46). This places the improved mitochondrial function at a time when endurance performance is highest. Thus, based on our citrate synthase data, it is possible that robust muscle clock phase shifts in the early active phase runners drives a phase shift in circadian clock directed gene expression leading to an earlier time-of-day enhanced mitochondrial function to support enhance performance adaptations. It is important to note that in the present study we did not directly measure skeletal muscle circadian phase or rhythmic mitochondrial function, and thus these proposed mechanisms remain hypothetical and require direct experimental confirmation in future work.

This model is also consistent with the outcomes from Xin and colleagues, (10) which demonstrated that endurance exercise performance can be improved, without training, through shifting the muscle clock with time restricted feeding. In Xin et al., (10) the authors found that 3 weeks of time restricted feeding during the rest phase in female mice was sufficient to significantly enhance endurance performance in the early light/inactive phase (ZT2). They went on to show that these endurance effects of time restricted required an intact circadian clock and implicate shifted rhythms in mitochondrial and lipid metabolic genes for this outcome. There is much still to be tested, but this model implicates the circadian clock system as a modifier of endurance performance adaptations. We also note that the different substrate metabolism required during ZT13 vs. ZT22 training also likely contributed to our observation that the early active phase runners lost the same amount of body fat as the late active phase runners with no impact on lean mass, even though the absolute training volume was significantly lower.

Finally, our findings may hold importance for understanding a new factor that may contribute to the growing efforts focused on understanding the heterogeneity of exercise responses, a major challenge in both human and translational research (47–49). Our study design implemented tight controls including inbred mice and clearly defined exercise training intensities and durations allowing us to identify exercise timing and performance outcomes. We do not suggest that exercise timing will solely explain exercise heterogeneity, but we do suggest that exercise timing is a variable that has been an underappreciated, yet biologically relevant, source of variability in exercise outcomes and should be a consideration in experimental designs.

This study is not without limitations. A relatively small sample size, combined with only female mice were used, which limits the generalizability of these data across sexes. Mitochondrial function, biogenesis, and stress responses have been shown to exhibit sexual dimorphism in both physiological and pathological <u>contexts (50), raising</u> the possibility that male mice could respond differently to time-of-day training. Replication of these findings in males will be important to determine the extent to which the observed effects are sex-specific. Also, we used an inbred C57BL/6 strain of mice, which was part of our control, but it also needs to be noted as a limitation as the application across more diverse genetic backgrounds remains to be tested. We also acknowledge that the western blot analyses were conducted in quadriceps muscle rather than gastrocnemius. Nonetheless, these datasets are consistent with enzymatic and performance outcomes, providing complementary but preliminary evidence that early active phase training promotes modest mitochondrial, contractile, and metabolic adaptations. Additionally, the study focused solely on endurance-type exercise training; resistance or high-intensity interval training may yield different time-of-day responses. Finally, all testing was performed at each group’s specific training time. Thus, we do not know if or by how much performance at different times of day might be impacted. Nevertheless, these findings have significant implications for exercise scheduling in both animal studies and human training regimens, as well as considerations for exercise performance following significant athlete travel. Optimizing training time to align the intrinsic muscle circadian rhythms may enhance the effectiveness of exercise interventions aimed at improving performance or metabolic health. However, additional studies are needed to clarify the molecular mechanisms at play, and to determine whether similar effects are observed across sexes, exercise modalities, and in different circadian contexts.

In conclusion, we report a minimum of 6 weeks of early active phase at ZT13 endurance training confers superior adaptive benefits compared to late active phase at ZT22 run training in mice, despite lower absolute training volumes. The greater adaptation per unit of work in ZT13-trained mice highlights a potential efficiency advantage tied to the timing of training, rather than total volume. Surprisingly, we also found that the early active phase runners exhibited the same magnitude of fat mass loss with training at a lower absolute training volume. These results suggest that the timing of exercise can significantly influence training efficiency and physiological adaptation, potentially via circadian regulation of skeletal muscle metabolism. We hypothesize that this enhanced adaptation to early active phase training may be due to shifts in the phase of the muscle clock, allowing for better alignment of the muscle’s metabolic capacity with exercise demands. While exercise is known to exert a myriad of positive health effects, the concept that these benefits may be advanced, in part, through changes in circadian rhythmicity, remains under studied. There is still much more work to be done in preclinical models to mechanistically test the exercise-induced changes in muscle circadian clock phase, mitochondrial capacity, substrate metabolism and endurance performance. It is also important to test the translation of these findings as they hold significant implications for athletic performance, experimental exercise-research design, as well as restoring functional deficits in individuals with circadian clock disruption e.g., type 2 diabetes and aging (51–54).

## DATA AVAILABILITY

The data that support the findings of this study are available on request from the corresponding author.

## SUPPLEMENTAL MATERIAL

Supplemental Figures. S1-S5: DOI.10.6084/m9.figshare.29476850

## AUTHOR CONTRIBUTIONS

SJ Hesketh and KA Esser conceived and designed the research; SJ Hesketh, CM Douglas, X Zhang, CA Wolff ES Nowicki and CL Sexton performed the research and acquired the data. SJ Hesketh CA Wolff, and CL Sexton analyzed and interpreted the data. SJ Hesketh drafted the manuscript, and all authors were involved in editing and approved the final version.

